# Fine-scale maternal structure and high nuclear connectivity in the French populations of the Geoffroy’s Bat (*Myotis emarginatus*)

**DOI:** 10.64898/2026.10.05.756748

**Authors:** S Blineau, O Tournayre, P Chevret, S Jacquet, R Leblois, L Belateche, J Duhayer, M Leuchtmann, A Pirog, C Régis, C Kaerle, D Pontier

**Affiliations:** Laboratoire de Biométrie et Biologie Évolutive UMR5558, CNRS, Université Lyon 1, Villeurbanne, France; ISEM, Univ Montpellier, CNRS, IRD, Montpellier, France; Maladies Infectieuses et Vecteurs : Ecologie, Génétique, Evolution et contrôle (MIVEGEC), Université de Montpellier, CNRS, IRD, Montpellier, France; CBGP, INRAE, CIRAD, IRD, Montpellier SupAgro, Université de Montpellier, Montpellier, France; Nature Environnement 17, Surgères, France; ANTAGENE, Animal Genomics Laboratory, 6 allée du Levant, 69890 La Tour de Salvagny, Lyon, France

**Keywords:** Chiroptera, mitochondrial DNA, microsatellites, gene flow, female philopatry, phylogeography

## Abstract

The Geoffroy’s bat (*Myotis emarginatus*) is a sedentary species increasingly threatened by habitat alteration. Here, we assessed genetic diversity and population structure in 760 individuals from 18 French maternity colonies using 34 microsatellite markers and mitochondrial D-loop sequences. Mitochondrial data revealed subtle regional structure, consistent with female philopatry, and a broader phylogeographic split between Western Europe and the Greater Middle East, reflecting historical isolation. In contrast, microsatellites showed high nuclear diversity and minimal population structure across France, with weak isolation by distance and kin clustering, indicating extensive male-mediated gene flow. Moreover, demographic inference suggested population stability, consistent with both marker types. This nuclear homogeneity and demographic stability are strikingly similar to the patterns observed in *Rhinolophus ferrumequinum* populations sampled at the same colonies, suggesting that reproductive aggregation can maintain connectivity even in sedentary species. Overall, *M. emarginatus* functions as a largely connected population at the national scale, although localized maternal lineages suggest the presence of functional phylopatric units around key maternity sites that warrant protection. Future genome-wide studies could refine conservation strategies by integrating fine-scale female philopatry with male-mediated connectivity.

## I. INTRODUCTION

Global decline in biodiversity is driven by five main threats: habitat destruction and fragmentation, pollution, overexploitation, introduction of invasive species, and climate change (Hogue & Breon 2022 ; IPBES 2019). Extinction rates have increased dramatically over the past century, reaching levels that would have required hundreds to thousands of years under natural conditions, suggesting that biodiversity may be undergoing a sixth mass extinction (Cowie et al. 2022 ; Ceballos et al. 2015). According to the IUCN, 28% of assessed species are classified as threatened, including 26% of mammals, with a similar pattern in France where one-third of native mammal species are threatened or near-threatened.

Genetic diversity is critical for species persistence (Reed & Frankham, 2003; Hughes et al., 2008; Kardos et al., 2021), yet its integration into conservation strategies remains limited. Insufficient consideration of genetic risks allows inbreeding depression and genetic drift to contribute to species vulnerability, often before demographic declines become apparent (Reed & Frankham, 2003; Frankham, 2015). Loss of neutral genetic variation, as measured by microsatellites or mitochondrial DNA, may reflect demographic processes that can ultimately reduce adaptive potential and resilience to environmental change (Hughes et al., 2008), while adaptive genetic variation is increasingly recognized as key to long-term survival (Teixeira & Huber, 2021; Kardos et al., 2021). Thus, maintaining genetic diversity is essential to sustain resilience and population viability, especially for species with particular life history traits (e.g. low fecundity, small effective population size) (Allendorf et al., 2012).

Bats (Chiroptera) are particularly vulnerable to habitat loss and population fragmentation (Festa et al., 2023). Indeed, their longevity, low fecundity, high metabolic rates, and nocturnal activity make them sensitive to environmental and anthropogenic pressures, such as pesticide exposure, forestry and agricultural practices, building renovation, and climate change (Voigt & Kingston, 2016; Frick et al., 2020). Nearly half of bat species in France are currently classified as threatened or near-threatened, according to the French Red List (IUCN France, 2017). Their life cycle, involving seasonal hibernation and reproduction, and sex-based spatial segregation is complex (Voigt & Kingston, 2016), resulting in marked seasonal variation in connectivity and heterogeneous patterns of regional connectivity (Pinaud et al. 2025), with consequences for both demographic processes (Ronget et al. 2025) and genetic patterns (Kerth et al., 2003). Long lifespan and low reproductive output can reduce effective population size, thereby increasing susceptibility to genetic drift and inbreeding (Frankham 2005; Hoban et al. 2023). Moreover, behavioral traits such as female philopatry and seasonal swarming may further shape fine-scale genetic structure in ways that are not readily apparent from demographic data alone (Kerth et al. 2000). Because mitochondrial DNA is maternally inherited, female philopatry can result in stronger or contrasting spatial genetic structure in mitochondrial compared to biparentally inherited nuclear markers, depending on dispersal or colonisation processes (Kerth et al. 2002; Kerth and Petit 2005). Together, these ecological, behavioral, and demographic characteristics shape population connectivity and genetic structure, highlighting the importance of integrating genetic data into conservation strategies (Allendorf et al. 2010; Ripperger et al. 2014).

*Myotis emarginatus* is a medium-sized vesper bat distributed across southern and central Europe, North Africa, and parts of the Middle East (IUCN, 2021). It is a sedentary species, with breeding and hibernation roosts typically located within a 50 km radius (Fleming, 2019; Frantz et al., 2022), even though movements of over 100 km between sites have been reported (Figure S1). Mating occurs in September during swarming, when males gather near maternity sites (Parsons et al., 2003; Parsons & Jones, 2003). Furthermore, females are philopatric, meaning that they return to the same maternity roosts (i.e. roosts where *females gather to give birth in summer)* every year (Arthur and Lemaire, 2021; Spitzenberger & Weiss, 2020).

Previous genetic studies on M. emarginatus have primarily investigated its genetic diversity at broad geographic scales (Frantz et al. 2022, Uvizl & Benda 2022): across the species’ Western Palearctic range, mitochondrial data revealed several major phylogeographic lineages, whereas western European populations belong to a single lineage characterized by a star-like haplotype network (i.e. a few widespread haplotypes shared across distant regions and numerous low-frequency haplotypes with more restricted distributions). This pattern is consistent with a postglacial expansion followed by local diversification. However, because mitochondrial DNA is maternally inherited, broad-scale phylogenetic patterns may not fully capture contemporary fine-scale population structure associated with female philopatry. Analysing mitochondrial variation at the scale of maternity colonies may therefore reveal patterns of local female lineage that complement those inferred from biparentally inherited nuclear microsatellite markers.

In this study, we combined mitochondrial sequences and nuclear microsatellite markers to characterise the genetic diversity and population structure of *M. emarginatus* at two complementary spatial scales. Mitochondrial sequences were analysed within both a French and a broader European phylogeographic framework, whereas microsatellite markers were used to investigate contemporary genetic structure among maternity colonies in western France, where fine-scale sampling was available. Combining these markers allows to detect subtle patterns of genetic diversity that might be overlooked otherwise. Our objectives were to: (i) assess contemporary genetic structure among maternity colonies in western France, (ii) explore whether mitochondrial markers reveal patterns consistent with female philopatry, and (iii) place these fine-scale patterns within a broader demographic and phylogeographic context. By integrating historical and contemporary genetic information, this study improves our understanding of population connectivity in a cave-dwelling bat species and offers insights to inform conservation strategies across spatial scales.

## II. MATERIAL AND METHODS

### 1. Ethical statements

The French Ministry of Ecology, Environment and Sustainable development provided the authorization for *M. emarginatus* capture over the period 2015-2020 (approval no. C692660703 from the Departmental Direction of Population Protection (DDPP), Rhône, France). All methods were approved by the Muséum National d’Histoire Naturelle (MNHN) and the Société Française pour l’Étude et la Protection des Mammifères (SFEPM).

### 2. Sample collection

Bat captures were conducted at 18 maternity sites in France, during the parturition period (June -July) in 2016 and 2018 (Figure 1). Individuals were captured at dusk using a harp trap at the entrance of each site. Bats were placed in individual cloth bags, identified, measured, and weighed prior to release. Wing membrane biopsies (patagium) were collected using a 3-mm diameter biopsy punch (Worthington Wilmer & Barratt, 1996). Samples were preserved in 10 mL of 95% ethanol or dried with silica gel until DNA extraction (Puechmaille et al., 2011). In total, 761 adult individuals (631 females, 124 males, and six individuals of unknown sex) were captured.

**Figure 1.**
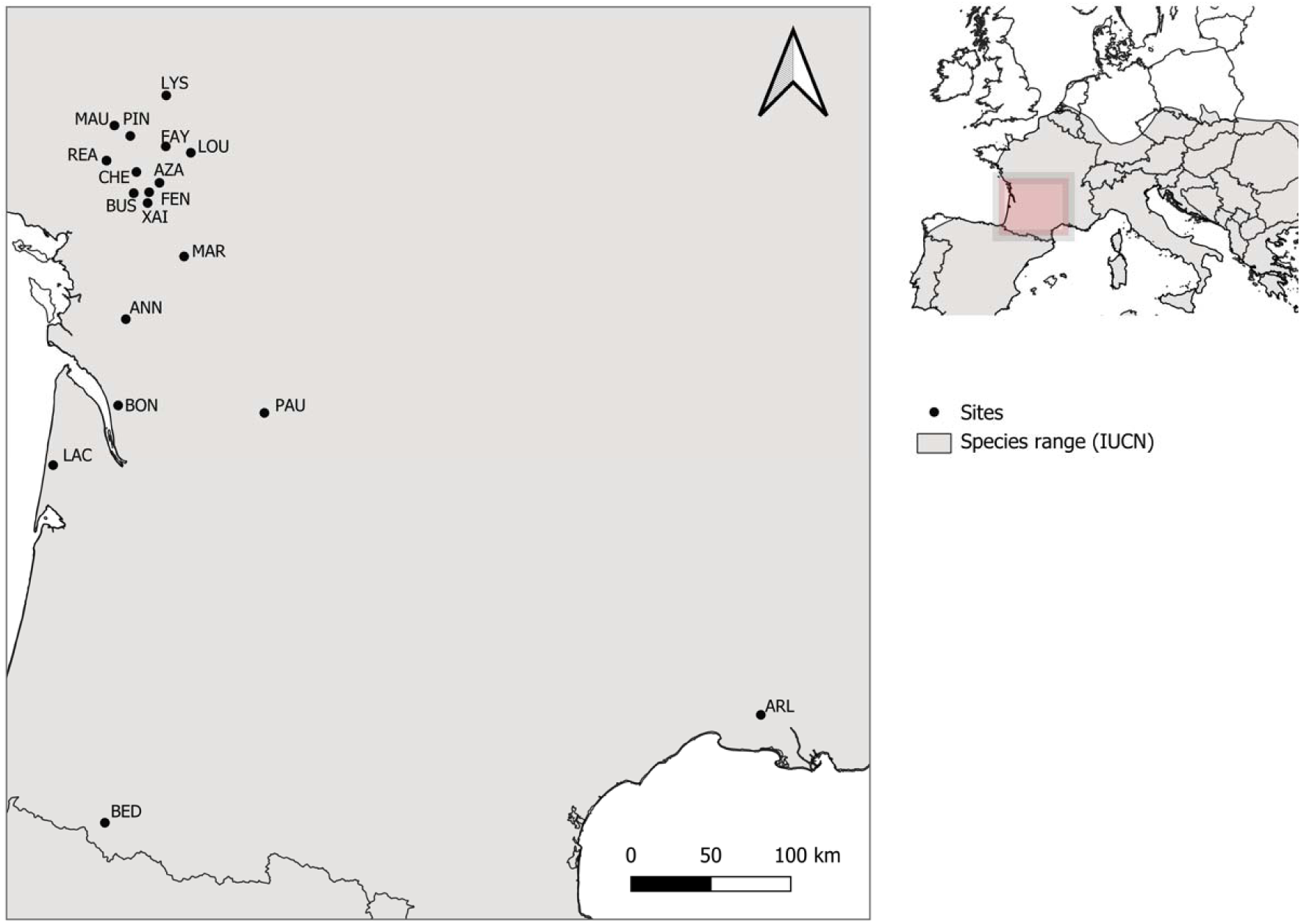
Location of the 18 *M. emarginatus* maternity colonies sampled in France. The overall study area is highlighted in red on the inset map of Europe). Sampled colonies are: Annepont (ANN), Arles (ARL), Azay-sur-Thouet (AZA), Bedous (BED), Faye l’Abesse (FAY), Fenious (FEN), Lacaneau (LAC), La Chapelle-Saint-Etienne (CHE), Le Busseau (BUS), Le Pin (PIN), Lys-le-Layon (LYS), Mauleon (MAU), Paussac et Saint-Vivien (PAU), Reaumur (REA), Saint-Bonnet-sur-Gironde (BON), Saint-Loup-Lamaire (LOU), Saint-Martin-les-Melle (MAR), et Xaintray (XAI). The species’ distribution range is shown in grey, based on IUCN data (2016; *The IUCN Red List of Threatened Species*, version 2025-2, accessed February 25, 2026).

### 3. Laboratory procedures

#### 3.1. gDNA extraction

Genomic DNA was extracted from the wing biopsy samples using the NucleoSpin® DNA RapidLyse kit (Macherey-Nagel), following the manufacturer’s protocol, with a final elution volume of 100 µL. Negative controls of extraction (i.e. only reagents) were simultaneously processed to detect potential contamination. The quality and quantity of DNA was visually checked on 1% agarose gels.

#### 3.2. Development and genotyping of microsatellites

Sequencing libraries were prepared using the Ion Xpress™ Plus Fragment Library Kit (Thermo Fisher Scientific; Waltham, USA) on 14 *M. emarginatus* samples. Libraries were subjected to double-sided size selection with SPRIselect Beads (Beckman-Coulter; Brea, USA), yielding DNA fragments ranging from 250 to 600 bp. Sequencing was performed on an Ion S5™ XL System using the Ion 530 Kit Chef (Thermo Fisher Scientific; Waltham, USA). All library preparation and sequencing steps followed the manufacturer’s recommendations to ensure consistent library quality. Microsatellite-enriched sequences were then identified and analysed using the QDD software pipeline (Meglécz et al. 2014). Primer pairs were designed based on expected amplicon size (≥150 bp), motif type (di-, tetranucleotide) and repeat number (≥ 10 for dinucleotides and ≥ 6 for tetranucleotides). Candidate loci were then screened for polymorphism by genotyping four individual samples and three pools of 10 individuals each on an ABI 3730 XL sequencer (Applied Biosystems, Foster City, CA, USA). After this selection steps, 29 primer pairs (Table S1) were retained for genotyping the complete dataset. These newly developed microsatellite loci were supplemented with eight microsatellite loci (Frantz et al., 2022) and two sex-linked microsatellite loci (Sánchez et al. 1996, Zarzoso-Lacoste et al. 2018). In total, 37 microsatellite markers were used to genotype the 761 *M. emarginatus* individuals (Table S1).

The 37 loci were arranged into three multiplex PCR reactions. All 761 samples were genotyped on 96-well microplates, along with three negative controls and three positive controls (samples previously analyzed and validated for DNA quality and genotyping success) per plate. Polymerase Chain Reactions (PCR) were performed in a final volume of 10 µL containing 5 µL of Mastermix Taq polymerase (Multiplex PCR Kit; Qiagen, Hilden, Germany), 1.30 µL , 1.62 µL or 2.12 µL of primer mix (A, B, and C, respectively, see Table S1) at concentrations ranging from 0.06 to 0.60 µM, and an average of 2 µL of genomic DNA. One primer from each pair was labelled with a fluorescent dye. The PCR thermal protocol consisted of an initial denaturation step at 95°C for 15 min, followed by eight touchdown cycles comprising 94°C for 30 s, annealing from 62°C to 55°C for 90 s (decreasing by 1°C per cycle), and 72°C for 30 s. These were followed by 29 cycles of 94°C for 30 s, 55°C for 90 s, and 72°C for 30 s, with a final extension step at 60°C for 30 min. PCR products were resolved using an ABI PRISM 3130 XL capillary sequencer (Applied Biosystems, Foster City, CA, USA) under denaturing conditions (HiHi^TM^ Formamide, Thermo Fisher Scientific) and an internal size marker. Electropherograms were analyzed using GeneMapper v.4.1 (Applied Biosystems).

Linkage disequilibrium among loci and departure from Hardy–Weinberg equilibrium were tested using Genepop v4.7.5 (Rousset, 2008). *P*-values were adjusted for multiple testing using the false discovery rate (FDR) procedure (Benjamini & Hochberg, 1995; Caldas de Castro & Singer, 2006).

#### 3.3. Mitochondrial sequencing

PCR amplification of the mitochondrial control region (DLoop) from the French sites was performed following the protocol described in Frantz et al. (2022) (i.e. a 342-bp fragment was amplified using the primers CR.HVII.F and CR.HVII.R). CytB amplification was performed using newly designed primers on a 989 bp fragment (Forward : 5’ ATGACCAACATYCGAAAAWC 3’, Reverse : 5’ GGYTGKCCTCCRATTC 3’). PCR reactions were carried out using the Q5 High-Fidelity DNA Polymerase from New England Biolabs (NEB), following the manufacturer’s instructions. Each reaction had a final volume of 25 μL, with 0.5 μM of each primer, and an annealing temperature of 58°C. Most colonies included eight to nine sequenced individuals. However, only four to five individuals were sequenced in neighbouring colonies because long-term monitoring using PIT-tagged females and automatic antennae (i.e. devices deployed at the entrances of maternity roosts) revealed sufficient movements of individuals among these geographically close colonies to expect limited mitochondrial differentiation between them. Thus, increasing the number of individuals sampled from each of these colonies was considered unlikely to provide substantial additional information. The quality of the PCR products was verified on an agarose gel and they were sequenced using the PCR primers by Microsynth (Vaux-en-Velin, France). In total, 110 D-loop sequences and 171 CytB sequences were obtained. Raw chromatograms were visually inspected, edited, and aligned using CLC Genomics Workbench (QIAGEN) prior to downstream analyses.

Finally, 1,031 D-Loop sequences were retrieved from GenBank (942 from Frantz et al., 2022 and 89 from Uvizl & Benda, 2022), including 62 sequences originating from France and 969 elsewhere in Europe. They were combined with the 110 sequences generated in this study, for a total of 1,141 sequences. All sequences were aligned using SeaView v4 (Gouy et al., 2010) with MUSCLE v3.8.31. Only the longest shared overlapping region was retained for downstream analyses (274 bp).

### 4. Analyses

#### 4.1. Microsatellite genetic analyses

##### Genetic diversity within colonies

For each locus, the proportion of missing data, the number of alleles, and the presence of potential null alleles were estimated using the adegenet R package (Jombart, 2008) and MICRO-CHECKER (Van Oosterhout et al., 2004). Within-colony genetic diversity was quantified by estimating allelic richness (*Ar*) and private allelic richness (*pA*), standardised to the minimum sample size using HP-RARE (Kalinowski, 2005). Observed (*Ho*) and expected heterozygosities (*He*) were calculated in Arlequin v3.5 (Excoffier et al., 2007). The inbreeding coefficient *F_IS_* (Weir & Cockerham, 1984) was estimated for each colony using Genepop v4.7.5.

##### Genetic differentiation and population structure

The global F*_ST_* was calculated using the adegenet and hierfstat (Goudet & Jombart 2022) R packages. Pairwise *F_ST_* values between colonies were calculated in Genepop v4.7.5 following Weir & Cockerham (1984), and exact G-tests of differentiation were performed. P-values were adjusted for multiple tests using the False Discovery Rate (FDR) procedure.

Population structure was further investigated using the Bayesian clustering approach implemented in STRUCTURE v2.3.4 (Pritchard et al., 2000), without prior information on population membership. Analyses were performed for *K* = 1-10, with 20 replicates per *K*. Each run included a burn-in of 1,000,000 iterations followed by 50,000 MCMC iterations. The most likely number of clusters was inferred using CLUMPAK (Kopelman et al., 2015), based on both Δ*K* values (Evanno et al., 2005) and log-likelihood scores. In addition, a Discriminant Analysis of Principal Components (DAPC) and associated genotype composition plot (compoplot) were performed using the adegenet (Jombart, 2008) R package.

The potential influence of kinship on population patterns was explored with pairwise relatedness coefficients (*r*) calculated within and among colonies using ML-RELATE (Kalinowski et al., 2006).

Finally, isolation by distance (IBD) analyses were performed in Genepop using the genetic distances between colonies (F_ST_/1−F_ST_ metric; Rousset, 1997) and the logarithm of the geographic distances, with significance of the regression slope and intercept assessed by bootstrapping over loci (default parameters). The correlation between the two matrices was then assessed using Mantel tests (10,000 permutations), with the adegenet R package to obtain a correlation coefficient that is no longer provided in Genepop output (both approaches yielded the same significance Mantel test results). The regressions were visualised using ggplot2 (Wickham, 2016).

##### Demographic inference

We inferred the demographic history of *M. emarginatus* using the maximum-likelihood coalescent-based approach implemented in MIGRAINE v0.6 (Leblois et al., 2014; Rousset et al., 2018). Because MIGRAINE assumes a strict stepwise mutation model (i.e. all alleles must be a multiple of the locus motif size), we excluded eight loci (ME17, ME22, ME24, ME28, Mema1, Mema5, Mema14, Mema32) that exhibited irregular repeat patterns incompatible with this assumption.

Analyses were first performed under the OnePopVarSize model, which assumes a single exponential change in population size starting *T* generations ago, with markers evolving under a generalized stepwise mutation model (GSM). This model estimates three scaled parameters: (i) the current scaled effective population size (θ = 4*N_e_µ*), (ii) the ancestral scaled effective population size (θ*anc* = 4*Nancµ*), and (iii) the scaled time since demographic change (*D* = *T* / 4*N_e_*), where *N_e_* and *Nanc* are the current and ancestral diploid effective population sizes, respectively, and µ is the mutation rate per generation. Evidence for demographic change was assessed using the ratio *Nratio* = *N_e_* / *Nanc*. Values < 1 indicate population contraction, whereas values > 1 indicate expansion. A demographic change was considered significant when the 95% confidence interval (CI) of *Nratio* did not include 1. Analyses were conducted using 400 parameter points, with 2,000 simulations per point, and six iterative refinements.

When no signal of demographic change was detected, we fitted the OnePop model (constant population size) to estimate θ, using 200 points, 200 simulations per point, and four iterations.

#### 4.2. mtDNA phylogeography

The final alignment of the France-wide dataset comprised 173 sequences of 274 bp (DLoop), including 63 sequences from previous studies (Frantz et al. 2022, Uvizl & Benda 2022). At the European scale, the final dataset included 1,141 sequences of 274 bp, of which 1,031 originated from published datasets (Frantz et al. 2022, Uvizl & Benda 2022). Haplotypes were identified separately for each dataset, and genetic diversity indices were calculated for each locality using DnaSP v6 (Rozas et al., 2017). Median-joining haplotype networks (Bandelt et al., 1999) were constructed in PopArt (Leigh & Bryant, 2015), and the geographic distributions of haplotypes was visualized at both the French and European scales. A phylogeny of the haplotypes was reconstructed using the Maximum Likelihood method implemented in PhyML (Guindon et al., 2010).

## III. RESULTS

### 1. Microsatellite analyses

All 37 loci were polymorphic, with an average of 7.59 alleles per locus (range: 2 -14). Three of the 37 microsatellite loci were excluded due to significant deviations from Hardy-Weinberg equilibrium (lower observed heterozygosity than expected), with one of these loci also showing evidence of null alleles. The remaining 34 loci exhibited no evidence of scoring errors, allele dropout, stuttering or linkage disequilibrium. Of the 761 individuals sampled, 760 were retained after excluding individuals with more than 50% of missing genotypes. The overall percentage of missing data in the final dataset was 1.87%.

#### Genetic diversity

Using the entire dataset, the Hardy-Weinberg (HW) test was significant (*p* = 0.0004, global *F_IS_* = 0.006), indicating a slight deficit of heterozygotes. However, the magnitude of this deviation was extremely low given the very small global Fis value. Local HW tests were non-significant for all colonies but LOU, FAY and AZA. *F_IS_*values were generally low across all sites (i.e. negligible deviations from HW equilibrium), ranging from -0.0073 to 0.0643. Expected (*He*) and observed (*Ho*) heterozygosity were similar across colonies, ranging from 0.54 to 0.60, indicating no evidence of high levels of inbreeding within colonies. Estimates of allelic richness (mean *Ar* = 4.31) and private allelic richness (mean *pA* = 0.04) revealed high levels of genetic diversity within colonies and homogeneity between colonies (Table 1).

**Table 1.** Mean genetic diversity parameters and standard deviations for all studied *M. emarginatus* maternity colonies in France. *F_is_* = inbreeding coefficient; *He* = expected heterozygosity; *Ho* = observed heterozygosity; *Ar* = allelic richness corrected for sample size; *pA* = private allelic richness corrected for sample size. No Fis values remained significant after FDR correction (adjusted α = 0.00007).

| *Locality* | *Locality code* | *N* | *F<sub>is</sub>* | *H<sub>e</sub>* | *H<sub>o</sub>* | *Ar* | *pA* |
| --- | --- | --- | --- | --- | --- | --- | --- |
| Lys-Haut-Layon | LYS | 33 | -0,0073 | 0,54948 | 0,55349 | 4,36 | 0,03 |
| Mauléon | MAU | 35 | 0,0226 | 0,57983 | 0,56715 | 4,25 | 0,03 |
| Le Pin | PIN | 47 | 0,0009 | 0,55788 | 0,55786 | 4,32 | 0,02 |
| Faye l'Abbesse | FAY | 63 | 0,0396 | 0,5644 | 0,5417 | 4,25 | 0,03 |
| Saint-Loup-Lamaire | LOU | 58 | 0,0244 | 0,55484 | 0,54159 | 4,24 | 0,02 |
| Réaumur | REA | 60 | -0,0004 | 0,55525 | 0,55568 | 4,33 | 0,02 |
| La Chapelle Saint-Etienne | CHE | 50 | -0,0028 | 0,55238 | 0,5541 | 4,23 | 0,02 |
| Azay-sur-Thouet | AZA | 36 | 0,0643 | 0,53624 | 0,50162 | 4,11 | 0,01 |
| Fenioux | FEN | 52 | 0,0103 | 0,55411 | 0,54855 | 4,27 | 0,01 |
| Le Busseau | BUS | 50 | 0,01 | 0,56712 | 0,56157 | 4,15 | 0,02 |
| Xaintray | XAI | 23 | -0,0341 | 0,53932 | 0,55697 | 4,18 | 0,01 |
| Saint-Martin-lès-Melle | MAR | 44 | 0,0028 | 0,57436 | 0,57266 | 4,33 | 0,02 |
| Annepont | ANN | 72 | 0,0115 | 0,57121 | 0,56476 | 4,32 | 0,04 |
| Saint-Bonnet-sur-Gironde | BON | 39 | -0,0134 | 0,56411 | 0,57162 | 4,35 | 0,08 |
| Paussac et Saint-Vivien | PAU | 30 | 0,0134 | 0,57326 | 0,56569 | 4,44 | 0,03 |
| Lacanau | LAC | 15 | 0,0134 | 0,58647 | 0,57872 | 4,38 | 0,05 |
| Arles | ARL | 26 | -0,0181 | 0,59141 | 0,6023 | 4,76 | 0,23 |
| Bedous | BED | 27 | -0,0205 | 0,57641 | 0,58754 | 4,34 | 0,06 |
| **Total** |  | 760 | / | / | / | / | / |
| **Mean** |  | / | 0,0065 | 0,56 | 0,56 | 4,31 | 0,04 |
| **Standard deviation** |  | / | 0,0229 | 0,02 | 0,02 | 0,14 | 0,05 |

#### Population structure in France

The global *F_ST_* across all colonies was very low (*F_ST_* = 0.006, 95% bootstrap CI: 0.004-0.007), indicating slight genetic differentiation. Similarly, pairwise *F_ST_* values were extremely low, ranging from -0.0015 (between MAU and PIN) to 0.0291 (between BON and XAI). While some significant pairwise comparisons involved relatively distant colonies (e.g. ARL, BED, LAC, PAU, BON) (Figure 1), most pairwise comparisons were not statistically significant (p > 0.05), further supporting weak differentiation among colonies. STRUCTURE analyses showed that the ΔK statistics (Evanno et al. 2005) and log-likelihood reached their maximum value at K = 3 (Figure 2). However, both DAPC and COMPOPLOT analyses revealed little to no genetic structuring among groups consistent with widespread admixture at the individual level (Figure S2), suggesting that the STRUCTURE signal correspond to weak differentiation in allele frequencies rather than distinct genetic clusters (Figure S2). Exploring alternative values of K did not substantially change this pattern.

**Figure 2.**
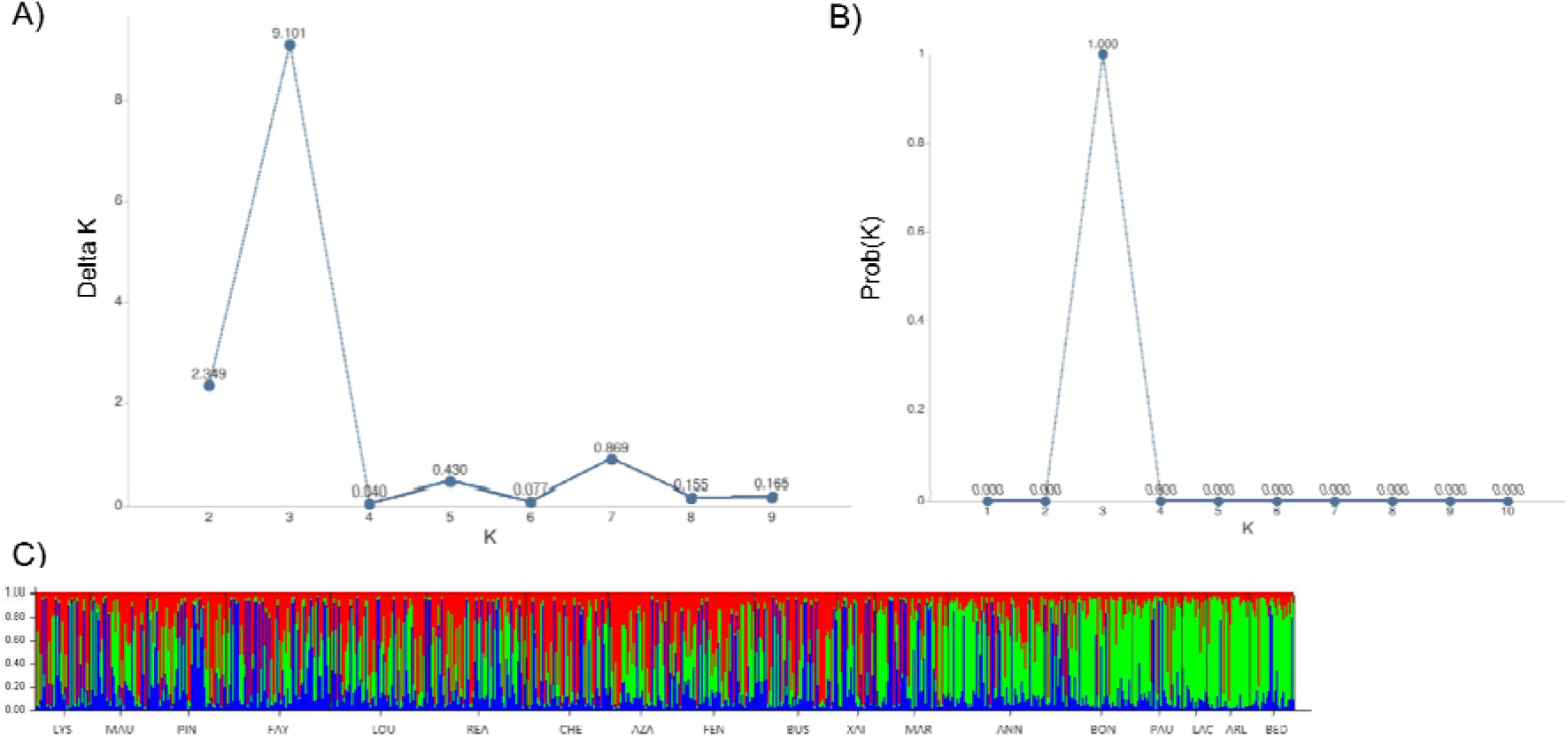
Population genetic structure inferred using STRUCTURE. (A) Evanno Δ*K* statistics used to identify the most likely number of genetic clusters, generated with from Clumpack ; B) Probability by K graph using median values of Ln(Pr Data), the k for which Pr(K=k); C) STRUCTURE plot of *M. emarginatus* for *K* = 3 clusters based on microsatellite data from 760 individuals. Each vertical bar represents an individual, and colors indicate the estimated membership probabilities to the three genetic clusters. The 18 sampled sub-populations are arranged from North to South along the X-axis.

Most individuals were unrelated both within colonies and across the full dataset (*r* = 0 for 49% of all inferred pairs, i.e. 141,315 out of 288,420 pairs) (Figure S3). The majority of pairs showing non-null relatedness (*r* values up to 0.69) involved geographically close colonies. Yet, rare occurrences of highly related individuals between distant colonies were observed: among the 114 pairs with r ≥ 0.5 (0.04% of all inferred pairs), seven pairs involved the Pyrenean colony BED and those from the core sampling area in western France (maximum *r* = 0.50). In contrast, no pair involving the furthest colony (ARL) reached *r* ≥ 0.5 (maximum *r* = 0.49), indicating a weaker and asymmetric connectivity with the rest of the sampled range (Figure S4). Moreover, the Mantel test revealed a significant positive correlation between genetic differentiation and log-transformed geographic distances across the full dataset (slope of the regression_GENEPOP_ = 0.005 [0.003,0.007] ; r*_MANTEL_* = 0.775, p-value_MANTEL_ < 0.0001) (Figure S5). Excluding the most distant colony (ARL), located on the Mediterranean coast, resulted in a smaller slope (a = 0.003 [0.002,0.005]) but still significant Mantel test (r*_MANTEL_*= 0.683, p-value_MANTEL_ < 0.0002) (Figure S5). The same pattern was observed when the second most distant colony (BED), located in the Pyrenean area, was also removed (a = 0.003 [0.002,0.005] ; r*_MANTEL_* = 0.650, p-value_MANTEL_ < 0.0003) (Figure S5).

Overall, the low *F_ST_*values, shallow isolation by distance slope, together with the lack of distinct genetic clusters, and low relatedness, indicate weak genetic structuring among maternity colonies consistent with substantial gene flow or connectivity among maternity colonies at the study area scale.

#### Demographic history

Inference under the OnePopVarSize model provided no evidence of demographic change in the *M. emarginatus* population (Table 2). Under the OnePop model, the scaled effective population size was estimated at θ = 1.696 [95% CI: 1.406-2.048], while the proportion of mutations following a generalized stepwise mutation model was estimated at *pGSM* = 0.327 [95% CI: 0.253-0.403].

**Table 2.** Demographic parameter estimates for *M. emarginatus* inferred using the OnePopVarSize model implemented in MIGRAINE. Estimates are based on 26 microsatellite loci and assume a generalized stepwise mutation model (GSM). θ and θ*anc* represent the current and ancestral scaled effective population sizes, respectively; θratio = θ/θ nc; *D* is the time since the demographic change; *g* is the time reported in generations; *µ* is the mutation rate per generation. Values in brackets indicate 95% confidence intervals.

| Parameters | Estimate | 95% CI |
| --- | --- | --- |
| pGSM | 0.331 | [ 0.259 - 0.401 ] |
| $\theta$ | 1.978 | [ 0.669 - 3.436 ] |
| $\theta_{anc}$ | 1.424 | [ 0.00111 - 202.5 ] |
| $\theta_{ratio}$ | 1.389 | [ 0.00931 - 1749 ] |
| Dg/2N | 0.0592 | [ 5.02e-05 - NA ] |
| Dgμ | 0.117 | [ 1.11e-05 - 1527 ] |

### 2. Spatial distribution of mitochondrial haplotypes

In our dataset, CytB sequences showed substantially low levels of diversity, with only three haplotypes differing by a single nucleotide mutation among the 180 individuals sampled in France. As a single haplotype was shared by all but six individuals, CytB data were not used in subsequent analyses.

In contrast, thirteen haplotypes were identified among the 173 D-Loop sequences sampled in France, and fifty-nine among the 1,141 D-Loop sequences sampled across the species’ distribution range (Figure 3A). The thirteen haplotypes identified in France were associated with haplotypes occurring elsewhere in Europe, Maghreb, and Turkey (Figure 3B). One haplotype (FR1) was widespread throughout Europe, Maghreb, and Turkey, but was absent from the Greater Middle East (North Levant, Holy Land, Oman, Iran, and Tajikistan) (Figures 3B). Several haplotypes detected in France (e.g., FR4 and FR6) were also found in other European countries, including Turkey, whereas others (e.g., FR5, FR8, and FR10) were restricted to western France. Haplotype FR9, common in Belgium and Luxembourg, was also detected in central and western France (Figure 3B), suggesting genetic connectivity among Western European populations. Overall, the distribution of haplotypes suggests variable levels of genetic connectivity across the species’ range.

**Figure 3.**
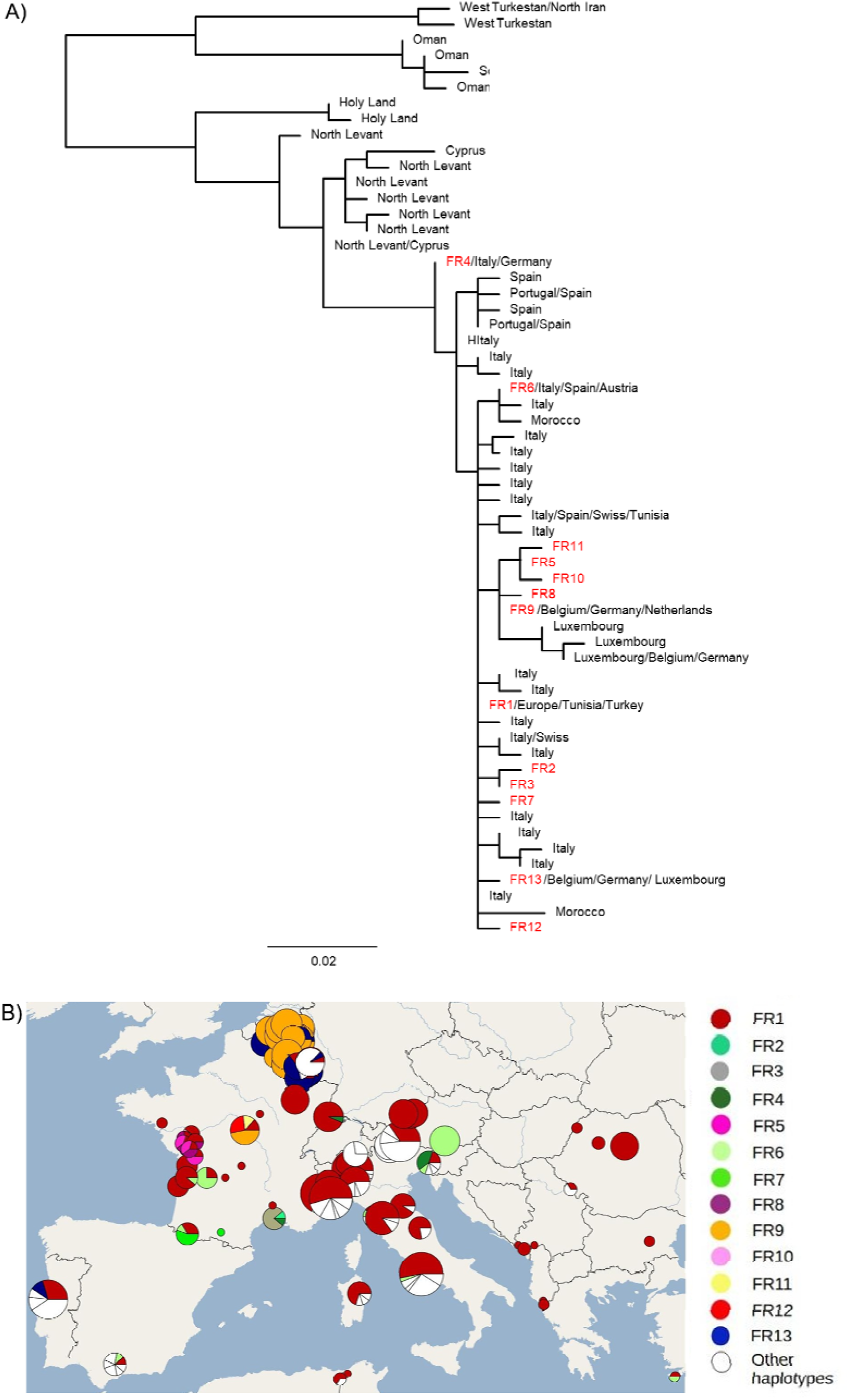
*M.emarginatus* haplotype diversity. A) Maximum likelihood phylogeny of the 59 haplotypes identified in the complete dataset. Haplotypes recorded in France are identified with the prefix “FR”. The country or geographic region is provided for each haplotype ; B) Pie charts representing haplotype frequencies within each colony, with pie size proportional to sample size and colored according to their geographic origin. For more clarity, only haplotypes recorded in France (« FR ») are colored.

The haplotypes identified in France differed from their closest relatives by a single mutation and from the central FR1 haplotype by a maximum of three mutations (Figure 4). The mitochondrial D-loop marker also revealed localised genetic differentiation in western, southern, and northern France, where several low-frequency haplotypes appeared to be geographically restricted (Figure 4).

**Figure 4.**
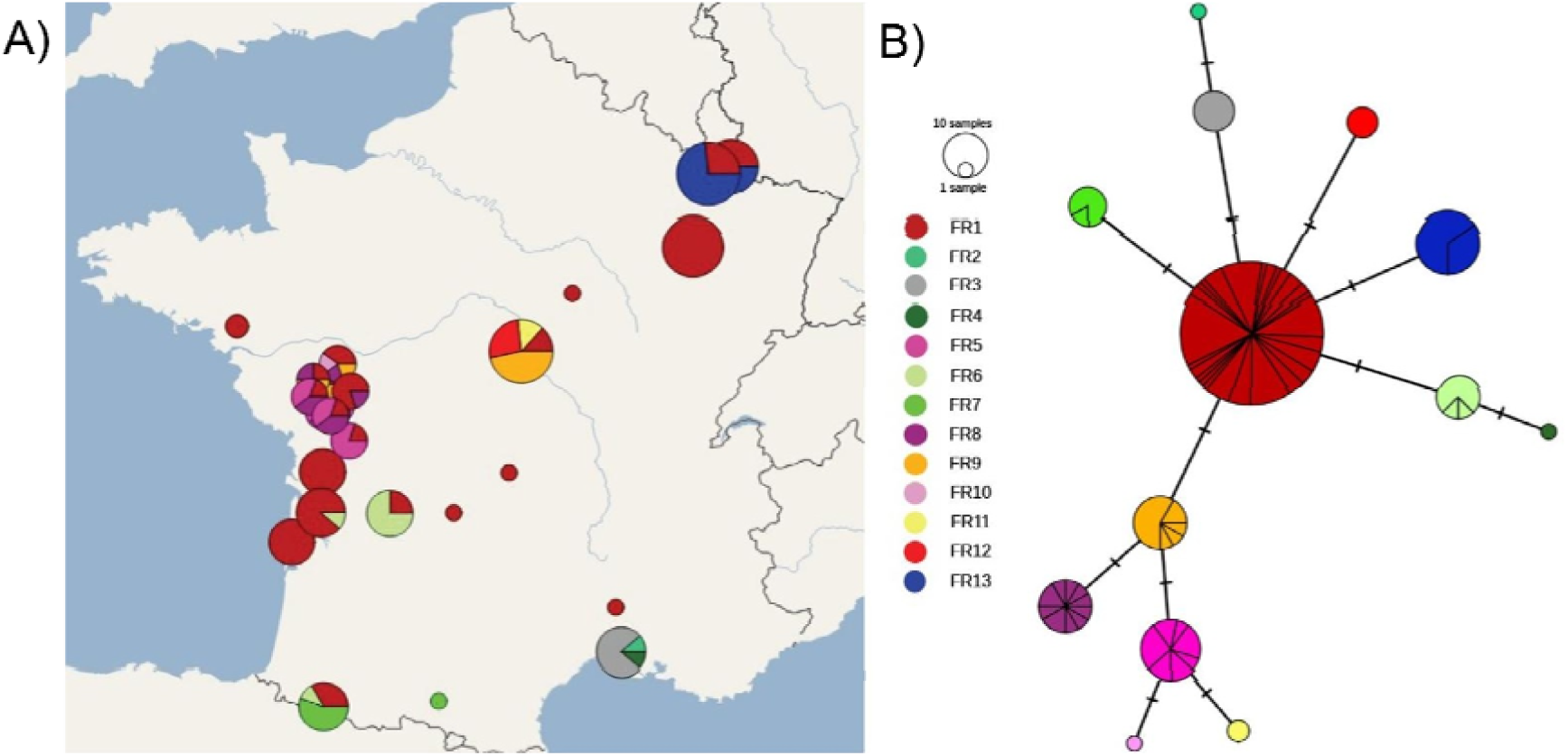
Distribution of mitochondrial D-loop haplotypes of *M. emarginatus* across France. Pie charts represent haplotype frequencies within each colony, with pie size proportional to sample size and colored sectors indicating haplotype proportions. (A) Geographic distribution of the 13 haplotypes identified in France; (B) Median-joining haplotype network at the French scale; circle size is proportional to the number of individuals carrying each haplotype, and hatch marks along branches indicate the number of mutational steps between haplotypes.

At a finer spatial scale, genetic diversity indices varied among samplingsites : haplotype diversity ranged from 0 (ANN, LAC) to 0.9 (i.e. four haplotypes identified among five sampled individuals ; BUS, LYS) with a mean of 0.555, and nucleotide diversity ranged from 0 to 0.005 with a mean of 0.003 (Table 3).

**Table 3.**
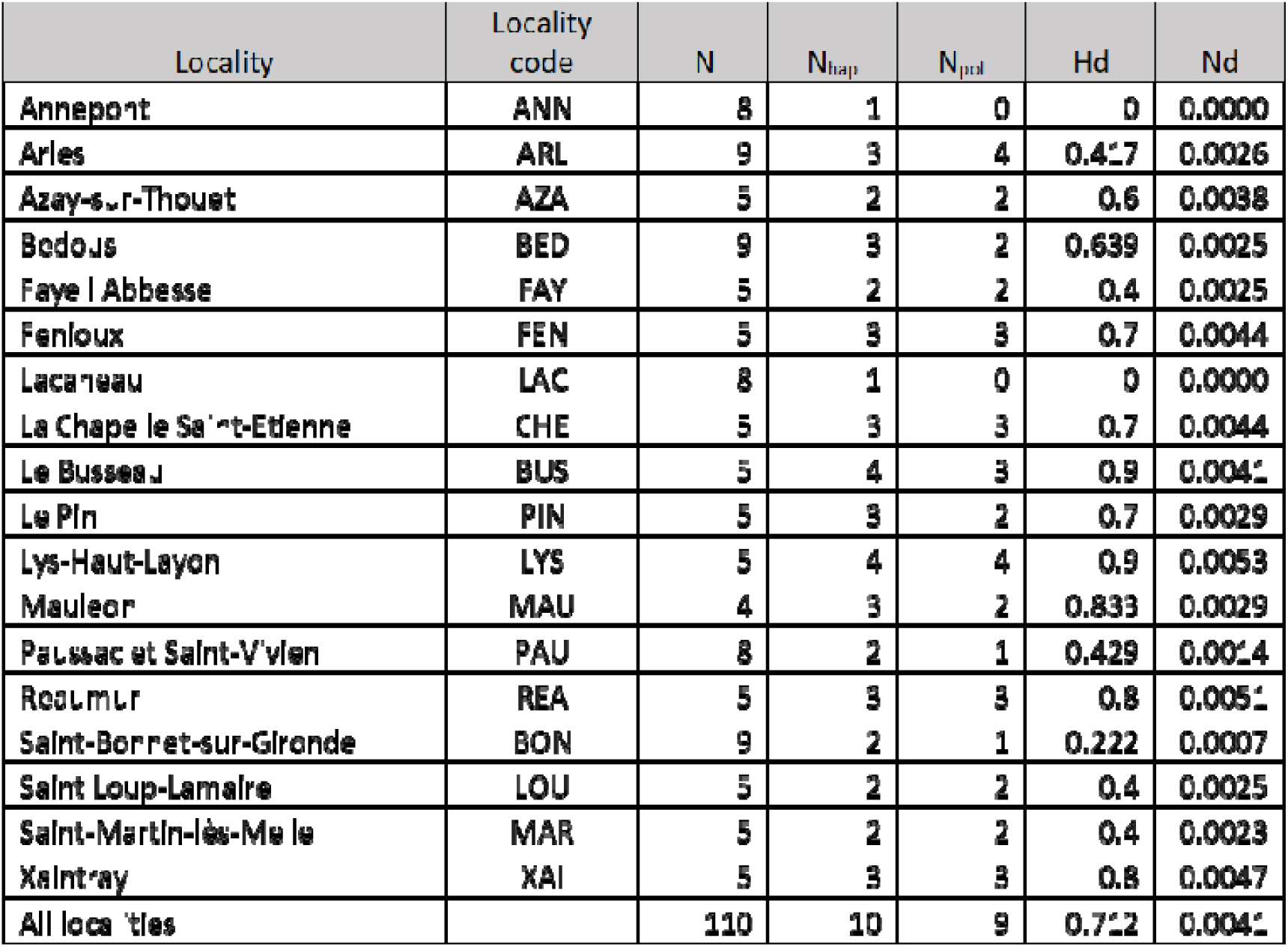
Mitochondrial genetic diversity (D-loop) in all studied *M. emarginatus* maternity colonies in France. *N_hap_* = number of haplotypes; *Npol* = number of polymorphic sites; *Hd* = Haplotype diversity; *Nd* = Nucleotide diversity.

## IV. DISCUSSION

Our results revealed a scale-dependent genetic structure shaped by the reproductive behaviour of this long-lived bat species. By combining mitochondrial and nuclear markers across a densely sampled area in western France and a broader European dataset, we found subtle mitochondrial differentiation among nearby maternity colonies, contrasting with high nuclear homogeneity at large scale. This discrepancy between marker types is consistent with female philopatry to maternity sites and nuclear gene flow inferred from nuclear markers mediated by late-summer swarming events. National-scale genetic homogeneity therefore likely reflects periodic mixing during these reproductive events rather than continuous large-scale dispersal. These findings highlight how life-history traits and mating systems can shape contemporary fine-scale genetic structure even within an otherwise broadly homogeneous European population.

### Continental phylogeographic continuity and local mitochondrial structure

At a broader geographical scale, the integration of our French samples into published European mitochondrial datasets revealed patterns consistent with the major mitochondrial lineages previously described at the continental level (Frantz et al. 2022, Uvizl & Benda 2022). Our results agree with the phylogeographic framework proposed by Frantz et al. (2022), who identified a dominant western Palearctic mitochondrial lineage characterised by a star-like haplotype network, including widely distributed haplotypes surrounded by geographically restricted variants. The presence of shared haplotypes across France and Europe may reflect common demographic history, including past population expansions and historical connectivity, although the retention of ancestral polymorphism may also contribute to this pattern. In contrast, the occurrence of more geographically restricted haplotypes suggests subsequent regional differentiation and local diversification processes. Similar combinations of widespread mitochondrial diversity and regional structuring have been reported by Uvizl & Benda (2022), supporting a scenario in which historical connectivity was maintained alongside the emergence of geographically structured variation. Overall, these patterns are compatible with a demographic history involving expansion from southern European refuge areas after glacial periods, followed by regional diversification of mitochondrial lineages (Frantz et al. 2022; Uvizl & Benda 2022).

Within France, spatial structuring was also observed. Several haplotypes (FR1, FR4, FR6) were shared among populations from France, Belgium, Austria, and Turkey. This broad distribution may reflect the persistence of ancestral mitochondrial variants following recent demographic expansion, historical connectivity among populations, or a combination of both processes.

### Female philopatry and population connectivity

The occurrence of haplotypes restricted to the northern, western, or southern clusters, with minimal sharing between regions, suggests fine-scale structuring among maternity colonies. Within France, these haplotypes differed from the dominant lineage by up to three mutations across ∼400 bp, indicating closely related maternal lineages. Their geographic restriction, together with the maternal inheritance of mitochondrial DNA, suggests limited female-mediated dispersal and strong fidelity to maternity sites. This pattern is congruent with those already documented in other temperate *Myotis* species, such as *M. lucifugus* (Dixon 2011) and *M. nattereri* (Scott et al. 2018), underscoring the importance of fine-scale sampling when assessing population structure in species with philopatric behaviour and strong reproductive aggregation.

In contrast, microsatellite analyses did not reveal any detectable genetic structure at the national scale. The slight but statistically significant global heterozygote deficit (*F_IS_* = 0.006) was considered biologically negligible, likely reflecting the pooling of weakly differentiated colonies rather than substantial inbreeding. This interpretation is consistent with the observed very low *F_ST_* values and the absence of discrete genetic clusters, supporting high connectivity among colonies. Although the STRUCTURE software identified an optimal K = 3, it likely reflects only subtle differences in allele frequencies rather than biologically meaningful population subdivision. Indeed, the DAPC, which does not rely on any of the population genetic assumptions underlying STRUCTURE (Pritchard et al., 2000; Jombart et al. 2010) showed no discrete assignment patterns, which was corroborated by the weak relatedness and isolation-by-distance patterns. Given the sedentary behavior of *M. emarginatus*, this nuclear homogeneity may not necessarily result from frequent long-distance dispersal. Instead, periodic reproductive gatherings such as late-summer swarming events may facilitate sufficient gene flow to homogenize allele frequencies across the landscape. Although the mechanisms maintaining this connectivity remain difficult to disentangle, long-term PIT-tag monitoring conducted at maternity roosts (Figure S1) provides independent ecological evidence that colonies are connected to some extent. Colonies sharing mitochondrial haplotypes could therefore be connected through common swarming sites, whereas more differentiated colonies could rely on distinct mating sites (Rivers et al., 2005). Interestingly, connectivity signals also suggested a potential spatial asymmetry: a few pairs of highly related individuals (r ≥ 0.5) linked the Pyrenean colony BED to colonies in the core western France sampling area, whereas no such pairs involving the Mediterranean colony ARL was observed (Figure S3). Although based on a small number of observations, this pattern is suggestive of potentially weaker or more geographically restricted connectivity with ARL, and should be further investigated in the future with denser sampling. Such asymmetry was also observed in the isolation by distance analysis. Excluding the Mediterranean colony reduced the regression slope by about 40% (from 0.005 to 0.003). However, the Mantel test remained highly significant, indicating that isolation-by-distance persisted at a relatively smaller scale. In contrast, excluding the Pyrenean colony BED did not reduce the slope any further, suggesting that BED’s genetic differentiation from other colonies scales with distance in the same way than the rest of the dataset. This is consistent with the relatedness analysis, which showed that BED, despite its geographic distance, remains connected to colonies in the core western France area.

The contrast between maternally inherited mitochondrial markers and biparentally inherited nuclear markers suggests that sex-biased dispersal and reproductive behavior may play a key role in maintaining connectivity at the national scale. A similar pattern has been reported in *Rhinolophus ferrumequinum*, a phylogenetically distant species sharing maternity and hibernation sites with *M. emarginatus,* and that was sampled from the same maternity colonies (Tournayre et al. 2019). Despite strong philopatry of both sexes to summer and winter roosts (Ronget et al. 2025, Rossiter et al. 2000), *R. ferrumequinum* also exhibits high nuclear homogeneity at the national scale, with weak IBD (slope of the regression within France = 0.006 [0.004–0.009]; pMantel < 0.001) and a clear majority of unrelated individuals (i.e. peak at *r* = 0) alongside some high relatedness values (*r* > 0.5) between individuals from both close and distant colonies (Tournayre et al. 2019). These patterns may result from mating outside natal colonies during autumn at shared transit or hibernation sites, where even relatively limited dispersal could be sufficient to maintain gene flow among colonies (Ronget et al. 2025). Altogether, these convergent patterns across two distantly related species suggest that in sedentary cave-dwelling bats, localised maternal lineages can coexist with high nuclear connectivity. More generally, they highlight the importance of considering both female site fidelity and male-mediated gene flow when delineating conservation units.

### Demographic stability

Demographic inference using MIGRAINE did not reveal any signal of demographic decline or expansion, supporting a scenario of long-term population stability. Some limitations of this approach, such as the lag in detecting recent demographic events, may have reduced its ability to detect recent demographic changes. The demographic analyses suggested a relatively large effective population size in *M. emarginatus*, with an estimated θ value of 1.696 (95% CI: 1.406–2.048). However, converting θ into an absolute estimate of *Ne* requires assuming a mutation rate, which remains uncertain for microsatellite markers in this species. Mutation rates can vary substantially among and within species, yet any uncertainty in the assumed mutation rate is directly propagated into estimates of *Ne* (Waples, 2010). Therefore, such conversions should be interpreted cautiously. A comparable absence of demographic decline and genetic structure has also been reported for *R. ferrumequinum* in the same region and maternity colonies (Tournayre et al., 2019), reinforcing the hypothesis that reproductive aggregation and inter-colony connectivity may contribute to maintaining effective population size in species exhibiting strong roost fidelity.

### Broader conservation implications

Rather than indicating genetic erosion, our results suggest that *M. emarginatus* currently retains sufficient connectivity to maintain neutral genetic diversity at the national scale. This implies that French colonies are unlikely to function as isolated genetic management units, but instead form part of a broader interconnected network. However, the presence of localized mitochondrial differentiation indicates that maternity colonies may still represent partially distinct maternal units. Protecting these key reproductive sites therefore remains essential for preserving fine-scale lineage diversity that underpins evolutionary potential of this species.

The similar patterns observed in the greater horseshoe bat *R. ferrumequinum*, which shares the same maternity and hibernation sites, further supports the idea that maintaining functional networks of roosts may benefit multiple species simultaneously. Conservation strategies should therefore prioritize landscape-scale connectivity among maternity and hibernation sites, rather than focusing solely on individual colonies.

Although our conclusions are based on neutral markers, genome-wide approaches integrating SNP data, ecological monitoring, and landscape analyses would help assess adaptive variation and finer-scale genetic structuring potentially organized around reproductive aggregation sites. Expanding sampling to underrepresented regions such as central France or the Mediterranean island (Corsica) would also help determine whether additional localized maternal lineages or reproductively connected colony clusters remain undetected. Integrating genomic and ecological data will be essential to refine conservation strategies while preserving the colony networks that sustain long-term demographic and genetic resilience.

## Author contributions

**Sterenn Blineau :** Data curation (lead), formal analysis (equal), investigation (equal), methodology (equal), visualization (lead), writing – original draft (equal). **Orianne Tournayre**: formal analysis (equal), investigation (equal), methodology (equal), visualization (equal), writing – review and editing (equal). **Pascale Chevret** : conceptualization (equal), investigation (equal), formal analysis (equal), methodology (equal), visualization (equal), supervision (equal), writing – original draft (equal). **Stéphanie Jacquet:** conceptualization (equal), investigation (equal), formal analysis (equal), methodology (equal), supervision (equal), writing – original draft (equal). **Raphael Leblois**: methodology (equal), formal analysis (equal), writing – review and editing. **Lilia Belateche**: methodology (equal), formal analysis (equal), writing – review and editing., **Jeanne Duhayer**: data curation (equal), writing – review and editing. **Maxime Leuchtmann :** data curation (equal), writing – review and editing, **Agathe Pirog :** data curation (equal), writing – review and editing. **Corinne Régis :** data curation (equal), writing – review and editing. **Cecile Kaerle:** data curation (equal), writing – review and editing. **Dominique Pontier**: conceptualization (equal), funding acquisition (lead), investigation (equal), methodology (equal), project administration (lead), supervision (equal), validation (equal), writing – original draft (equal).

## Acknowledgements

We are indebted to Jean-Baptiste Pons, Ondine Filippi-Codaccioni and to all of our collaborators in the naturalist associations who helped us to bat sampling: France Nature Environnement Nouvelle-Aquitaine, Poitou-Charentes Nature, Deux-Sèvres Nature-Environnement, Nature-Environnement 17, LPO France, LPO DT Aquitaine, Charente Nature, Vienne Nature, Groupe Mammalogique et Herpétologique du Limousin, Groupe Chiroptères Aquitaine, Cistude Nature, Conservatoire des Espaces Naturels de Nouvelle-Aquitaine, Société d’Étude et de Protection de la Nature en Lot-et-Garonne, Groupe Chiroptères des Pays-de-la-Loire, Groupe Chiroptères de Midi-Pyrénées, CPIE Vallée de la Sarthe et du Loir, LPO Vendée, LPO Anjou, LPO Centre Val de Loire, Parc National des Pyrénées, Parc Naturel Régional du Marais Poitevin, Comité départemental de spéléologie du Lot, Association des Naturalistes d’Ariège, and all the numerous volunteers that contributed to fieldwork. We also want to thank Alain C. Frantz for providing detailed files for his sequences, including localization information.

## Funding

Financial support was received from the LABEX ECOFECT (ANR-11-LABX-0048) of Université de Lyon, within the program “Investissements d’Avenir” (ANR-11-IDEX-0007) operated by the French National Research Agency (ANR), the RESPOND program of the Université de Lyon (UDL), Nouvelle-Aquitaine region, Nouvelle-Aquitaine DREAL, and European Regional Development Fund.

## Conflict of interest

The authors declare no conflicts of interest.

## Data availability statement

The raw data supporting the main results of the study are in the EMBL repository (accession number: OZ488857-OZ489138) and FigShare (microsatellites: https://figshare.com/s/c9c5f394924e9cc1e8af?file=67729755)

## SUPPLEMENTARY MATERIAL

**Figure S1.**
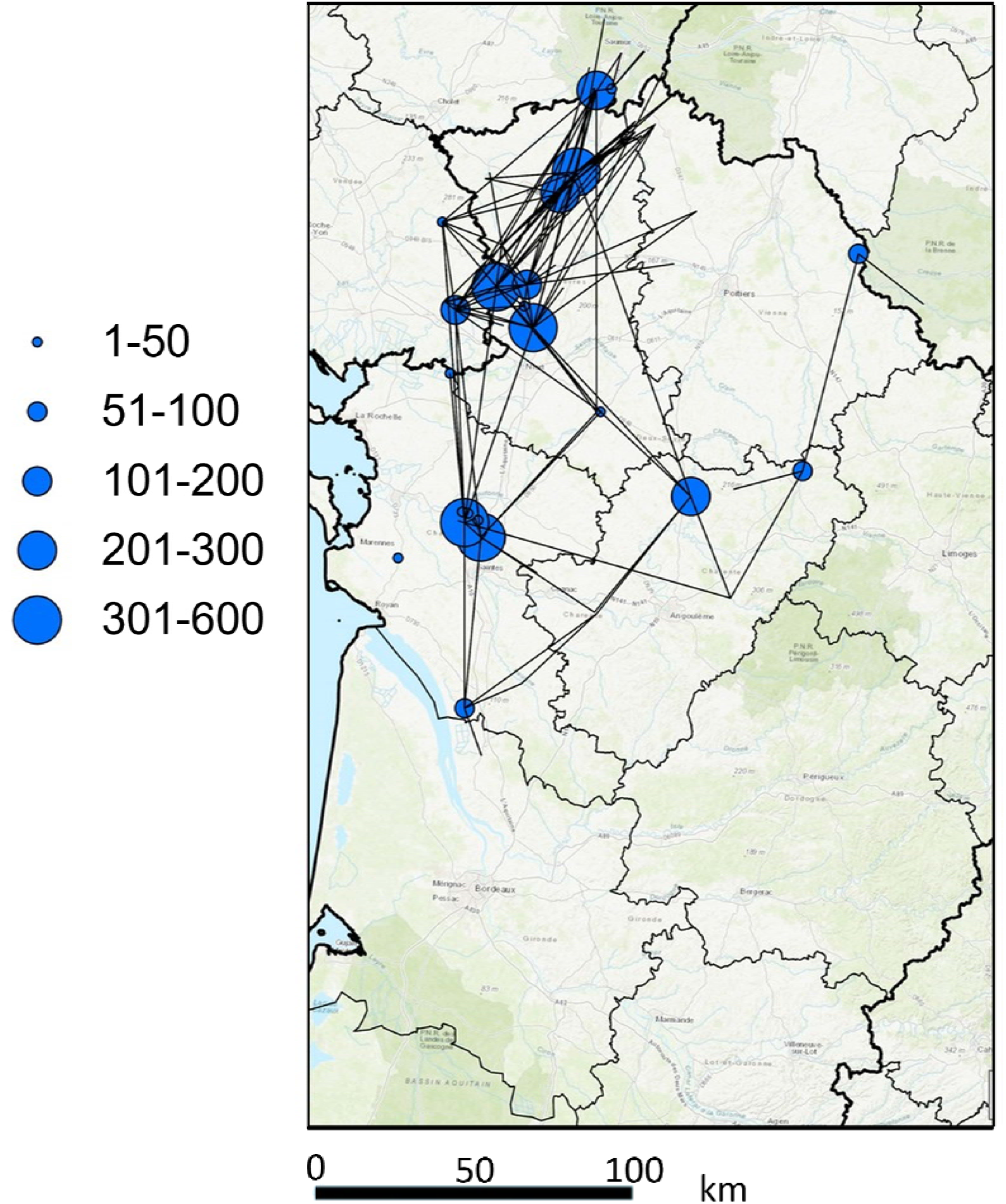
Movements (lines) of *M. emarginatus* individuals between roosts in western France. Blue dots indicate sites where individuals were PIT-tagged (mainly maternity colonies), with dot size proportional to the number of individuals tagged since 2019. Lines connect sites where individuals were subsequently detected or recaptured, including maternity colonies as well as transit and hibernation sites monitored via fixed or handheld antenna readers; these are not shown as dots to avoid overloading the map. Data source: France Nature Environnement Nouvelle-Aquitaine (2026), *Programme Chiroptères Cavernicoles Prioritaires en Nouvelle-Aquitaine* (CCPNA) and *Programme Université du pays Basque* (FNE Nouvelle-Aquitaine). Map layer: Geoservices IGN / ArcGIS Pro.

**Figure S2.**
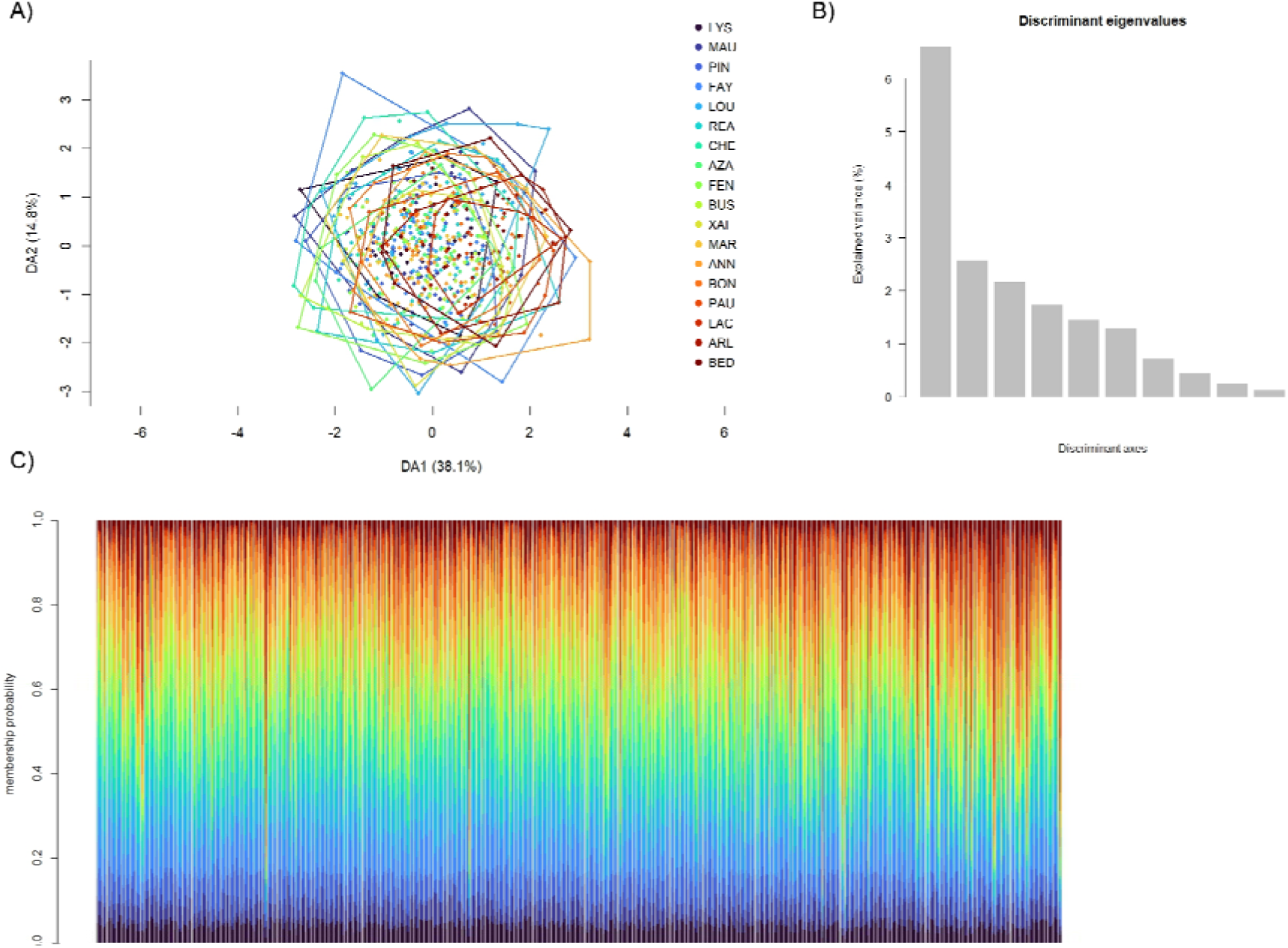
Population genetic structure inferred from a discriminant analysis of principal components (DAPC). (A) DAPC scatterplot. (B) Associated discriminating eigenvalues. (C) COMPOPLOT displaying individual admixture proportions. Colors indicate sampling sites.

**Figure S3:**
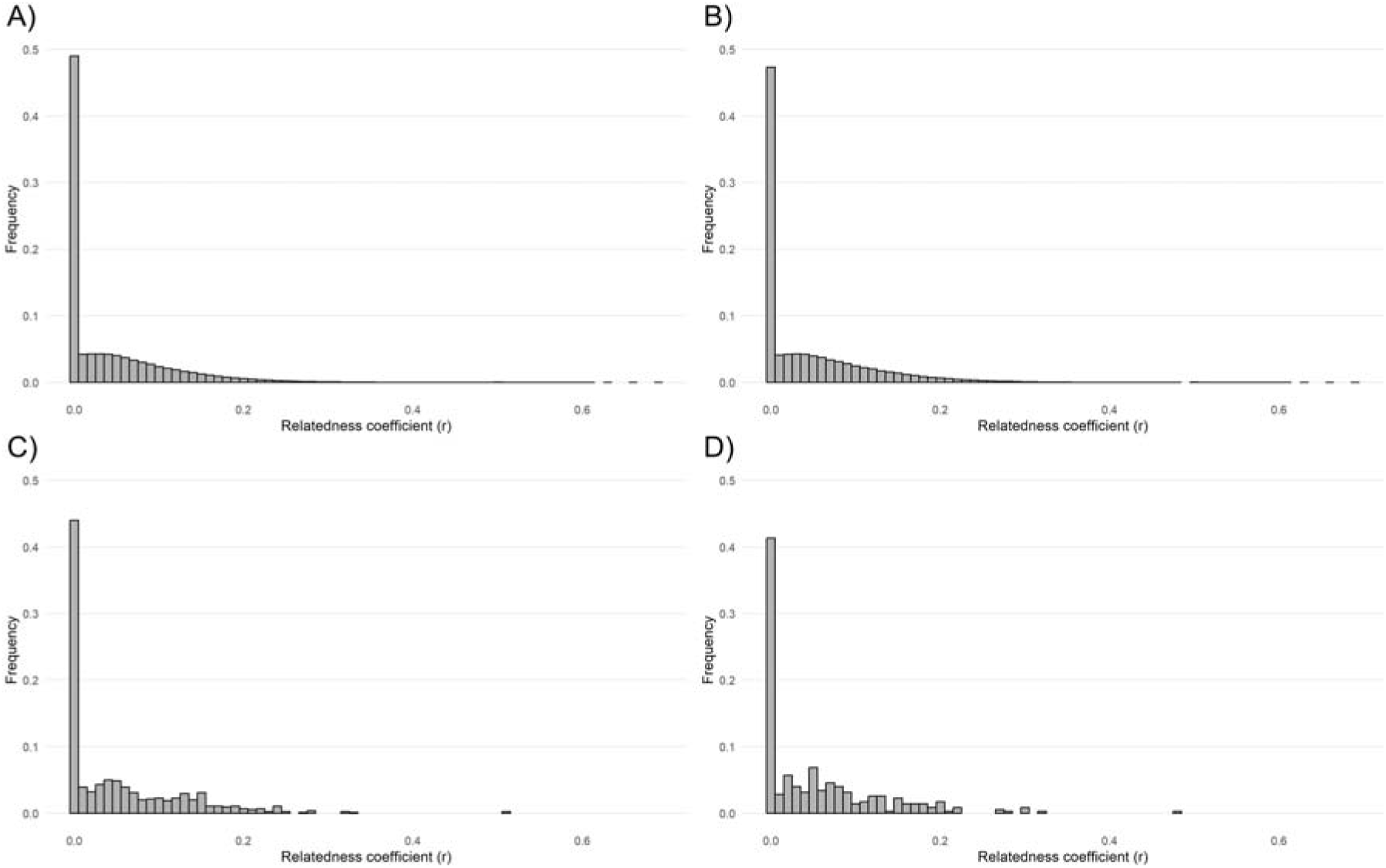
Distributions of the relatedness coefficient r across A) all sites, B) sites within the geographic western cluster (from LYS to MAR), C) the BON site only, and D) the BED site only. These two sites were randomly chosen.

**Figure S4.**
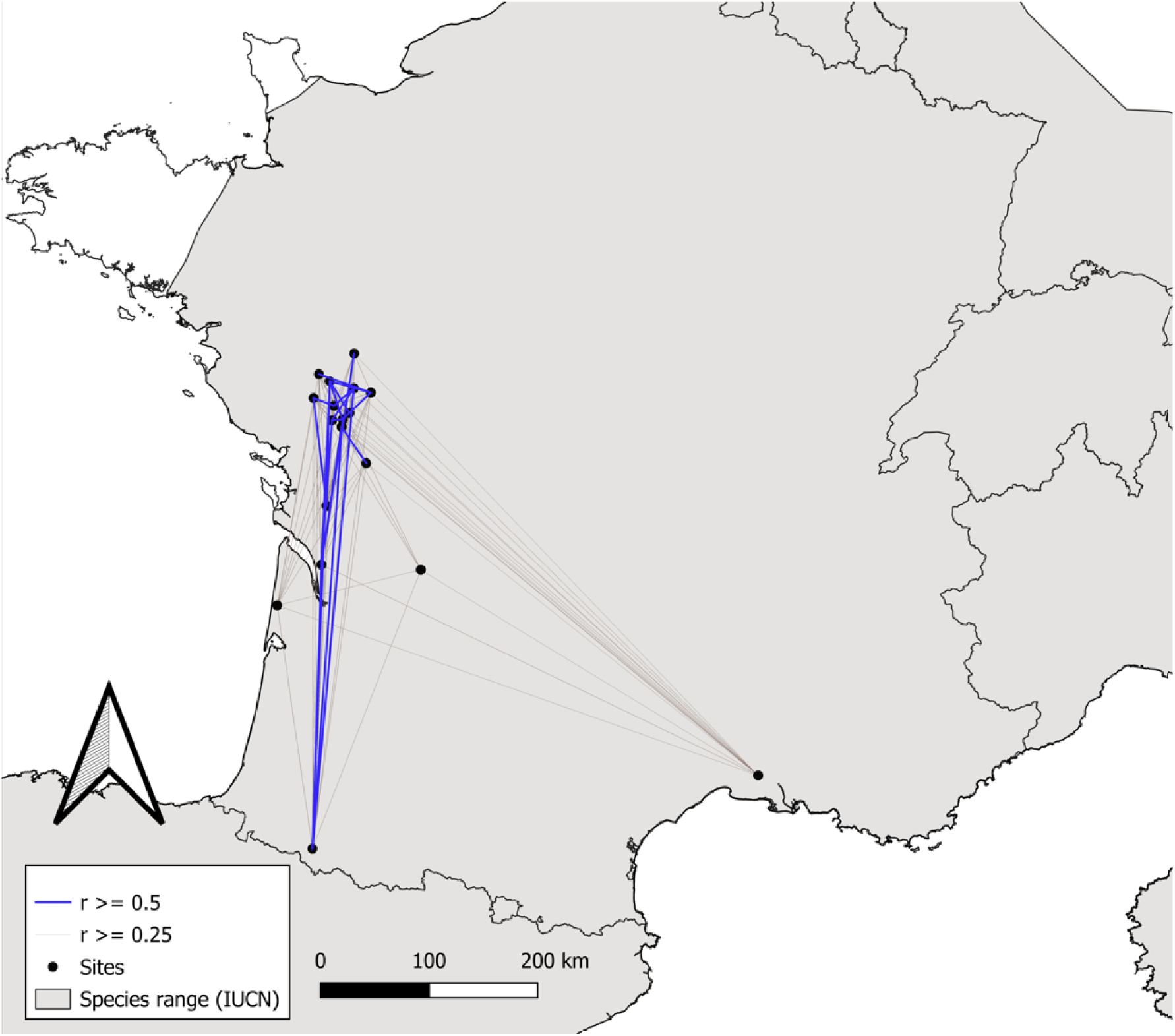
Geographic distribution of the highest pairwise relatedness coefficients (r ≥ 0.5 in blue and 0.25 ≤ r < 0.5 in grey) between *M. emarginatus* colonies in France. Note that the majority of pairs of individuals were unrelated (*r* = 0) but are not shown on this figure for clarity.

**Figure S5.**
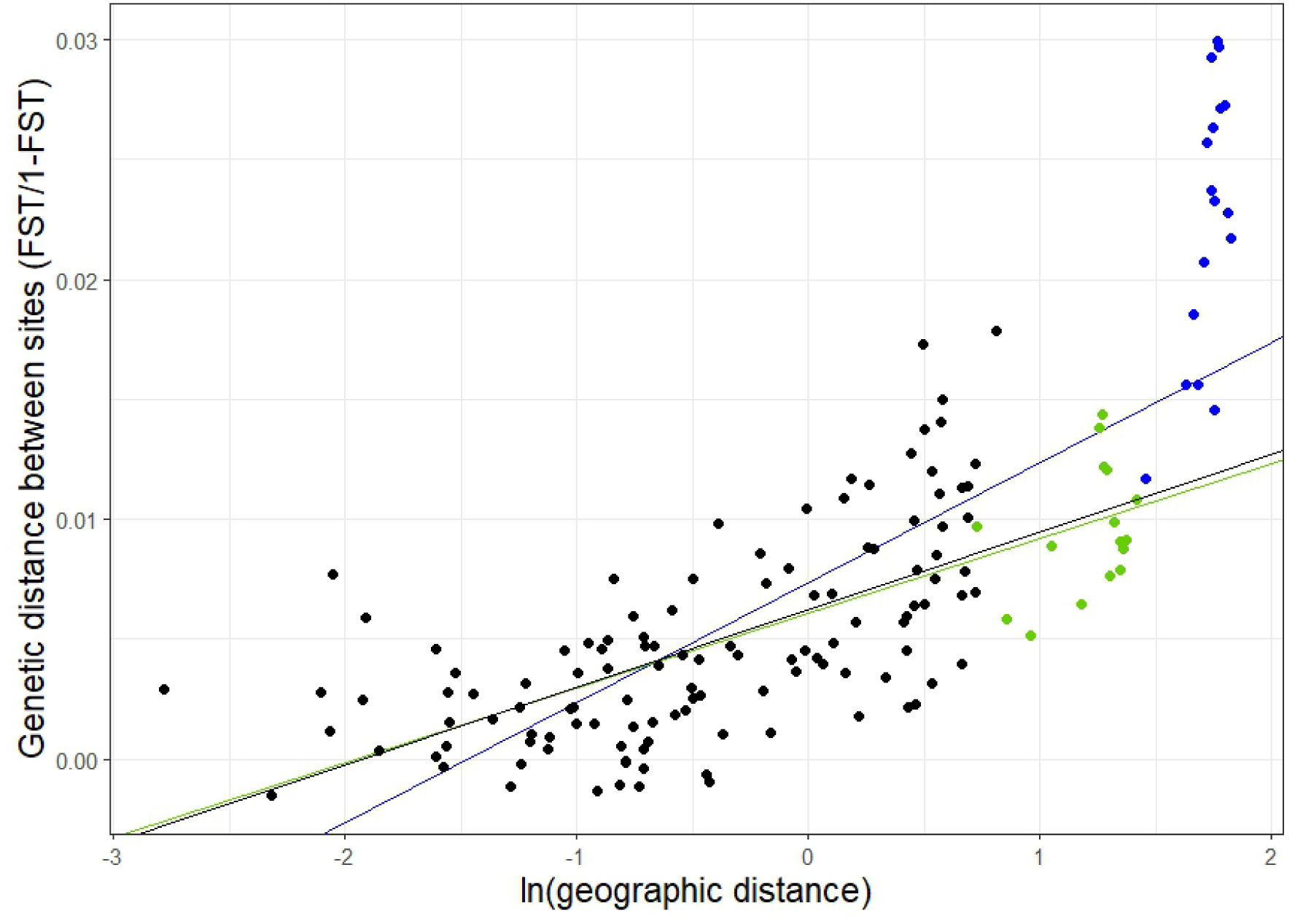
Plots of the regressions obtained in the isolation by distance analyses. Blue dots indicate pairs involving the Mediterranean ARL colony, and green dots indicate pairs involving the Pyrenean BED colony. These are the two most distance colonies. The blue line shows the regression including all colonies, while the almost overlapping green and black lines show the regressions after excluding ARL, and after excluding both ARL and BED, respectively.

**Table S1.**
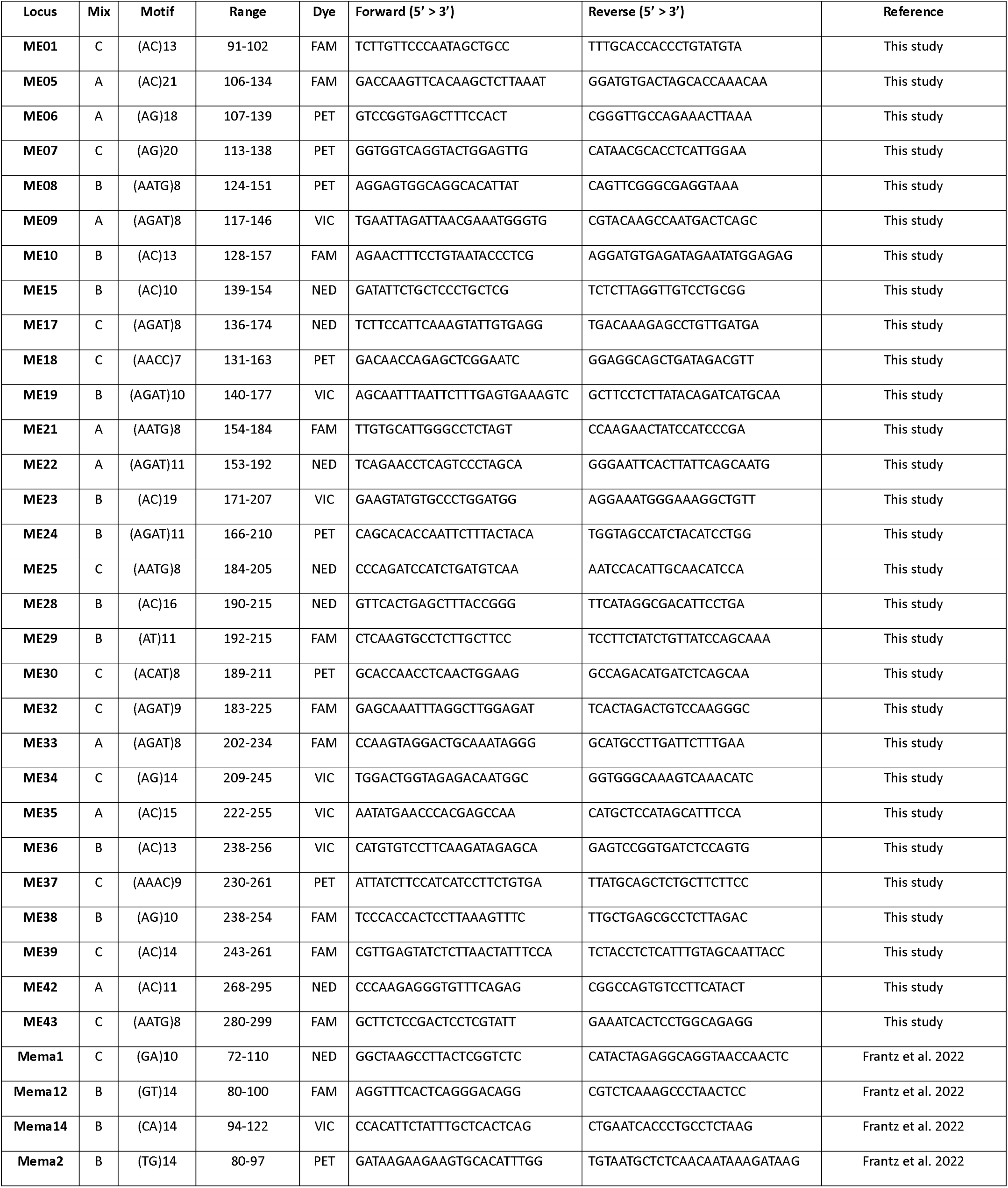
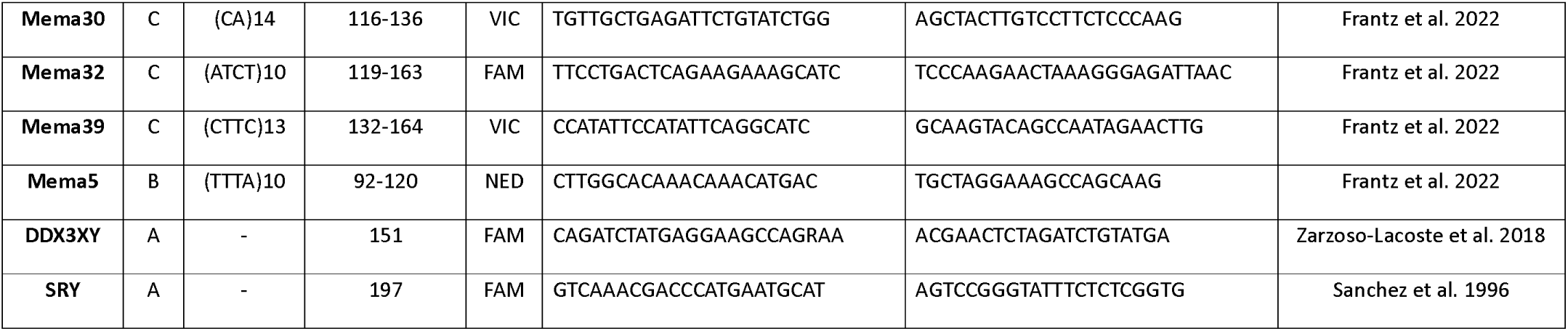
Information on the primer pairs used in this study.

## Notes

### Competing Interest Statement

The authors have declared no competing interest.

